# Single-cell genomics reveals opportunistic *Enterobacterales* carrying putative cationic antimicrobial peptide resistance genes in red crown rot-affected soybean rhizoplanes

**DOI:** 10.64898/2026.04.07.716964

**Authors:** Takeru Ochi, Yohei Nishikawa, Masako Kifushi, Takashi Sato, Haruko Takeyama

## Abstract

1.

**Abstract:** Soybean red crown rot, caused by the soil-borne fungus *Calonectria ilicicola*, causes substantial yield losses, but the response of the root-associated bacterial microbiome remains poorly understood. Here, we combined 16S rRNA gene sequencing, shotgun metagenomics, and single-cell genomics to characterize bacterial communities in soybean root-associated soils.

16S rRNA gene sequencing showed that diseased plants had rhizosphere and, more strikingly, rhizoplane microbiomes distinct from those of healthy plants, often with increased *Enterobacterales*. Shotgun metagenomics further revealed enrichment of genes associated with antibiotic resistance, particularly cationic antimicrobial peptide resistance, in diseased rhizoplane samples.

Single-cell genomics recovered seven nonredundant *Enterobacterales* genomes and showed that plant pathogenicity-related genes were broadly distributed across these lineages. In contrast, *dlt* genes, which are associated with cationic antimicrobial peptide resistance, were detected only in the *Enterobacterales* lineages enriched in diseased rhizoplane soils. These results support a model in which soybean red crown rot is accompanied by microbiome restructuring and opportunistic enrichment of specific *Enterobacterales* lineages carrying putative cationic antimicrobial peptide resistance genes. More broadly, this study highlights the value of strain-resolved single-cell genomics for linking disease-associated community shifts to specific bacterial traits.

**Importance:** Understanding crop disease requires resolving not only the primary pathogen but also the root-associated bacteria that respond to infection. Here, we used 16S rRNA gene sequencing, shotgun metagenomics, and single-cell genomics to examine the soybean rhizoplane microbiome under red crown rot. Diseased plants showed reproducible shifts in bacterial composition, including frequent enrichment of *Enterobacterales* and antimicrobial resistance-related functions. Strain-resolved genomes further revealed that the *Enterobacterales* lineages enriched in diseased rhizoplane soils specifically carried putative *dlt*-mediated resistance to cationic antimicrobial peptides, whereas general pathogenicity-related genes were broadly shared. These findings suggest that host defense-associated selection, rather than pathogenicity genes alone, may help shape disease-associated root microbiomes. This study demonstrates how single-cell genomics can uncover strain-level traits hidden within bulk community data and thereby clarify plant–pathogen–microbiome interactions.

## 2. Introduction

Soybean is one of the world’s most widely cultivated crops and a major source of plant-derived oil and protein for human consumption (1, 2). As global demand for soybean-derived products continues to increase, improving yield stability has become increasingly important (3–5). However, soybean production is substantially constrained by disease, with reported yield losses of 11.0% to 32.4% attributable to plant pathogens (6). Reducing the impact of soybean diseases is therefore essential for both food security and agricultural sustainability.

Soybean red crown rot (RCR), also referred to as black root rot, is an important soybean disease caused by the soil-borne fungus *Calonectria ilicicola* (7, 8). Although *C. ilicicola* can be detected by established methods such as microsclerotial isolation and quantitative PCR (9, 10), its detection alone does not fully explain field-level disease patterns. Disease incidence and severity vary among regions and years, and even within a single field, plants can differ markedly in symptom severity despite exposure to the same pathogen source. These observations suggest that additional factors, beyond the presence of *C. ilicicola* itself, may influence RCR development and severity (11).

Here, we considered root-associated bacteria as one such factor. Plants interact intimately with soil bacteria, and the rhizosphere and rhizoplane are well known to harbor host- and condition-specific microbial communities (12) . In plant disease systems, microbial interactions extend beyond the primary pathogen: co-occurring bacteria can respond to tissue damage, altered root exudation, and host immune activation, and may in turn influence disease progression (13–16). During disease, plants can alter root exudation and release antimicrobial peptides (AMPs) and other stress-associated compounds, thereby reshaping nearby microbial communities. However, the bacterial communities associated with soybean RCR have not been comprehensively characterized, and it remains unclear which bacterial lineages and functions are associated with the diseased root environment.

In this study, we analyzed the taxonomic and functional features of soybean root-associated bacterial communities using complementary genomic approaches. Bulk soils were collected at the sowing stage from multiple soybean fields in Akita Prefecture, Japan, and healthy and diseased soybean plants were collected from affected fields at the ripening stage. Soils were fractionated into bulk, rhizosphere, and rhizoplane compartments, followed by 16S rRNA gene sequencing and shotgun metagenomics to evaluate community composition and functional gene profiles. We further applied a microfluidic droplet-based single-cell genomics platform (SAG-gel) (17, 18) to recover strain-resolved bacterial genomes from rhizoplane soils. By integrating single-cell genome data with shotgun metagenomes, we identified bacterial lineages enriched in diseased rhizoplane soils and characterized their genomic features. Our results indicate that soybean RCR is associated with reproducible restructuring of the rhizoplane microbiome and enrichment of specific *Enterobacterales* lineages carrying putative CAMP resistance genes.

## 3. Result

### 3.1. A distinctive bacterial community dominated by *Enterobacterales* forms in the soybean rhizoplane affected by RCR

In June 2024, bulk soils were collected at the sowing stage from seven soybean fields (NS1 to NS7) in Akita Prefecture, Japan. During cultivation, localized poor growth suggestive of RCR was observed in field NS4. At the ripening stage, three diseased plants (replicates A to C) and three healthy plants (replicates A to C) were collected from NS4, and rhizosphere and rhizoplane soils were fractionated from each plant. All collected soil samples were analyzed by quantitative PCR using *C. ilicicola*-specific primers and by 16S rRNA gene sequencing targeting the V3–V4 region.

Quantitative PCR (qPCR) showed that *C. ilicicola* DNA in bulk soils collected at the sowing stage was below the detection limit in all seven fields (6.99 × 10^-1^ copies/µL of input DNA) (Fig. 1A). At the ripening stage, *C. ilicicola* DNA was also below the detection limit in all soil fractions (bulk, rhizosphere, and rhizoplane) from healthy soybean plants collected in NS4. In contrast, *C. ilicicola* DNA was detected in all soil fractions from diseased soybean plants collected in NS4, with values ranging from 5.46 × 10^1^ to 1.67 × 10^3^ copies per µg of soil DNA (Table S1). These results were consistent with the classification of the sampled plants as diseased or healthy. 16S rRNA gene sequencing showed that bulk soils collected at the sowing stage from NS4 were dominated by *Burkholderiales*, *Sphingomonadales*, and *Acidobacteriales*, all of which are commonly detected in agricultural soils (Fig. 1A) (19–21). This order-level composition was similar to that of the other six fields sampled in 2024 (Fig. S1A). At the ripening stage, bacterial composition in bulk soil remained broadly similar between healthy and diseased plants. In the rhizosphere and rhizoplane soils of healthy soybean plants, *Rhizobiales*, primarily *Bradyrhizobium*, were enriched, consistent with typical soybean root-associated communities (22). In diseased plants, *Rhizobiales* were also abundant, but *Enterobacterales* showed a marked increase in replicates A and B. In the rhizoplane, the relative abundance of *Enterobacterales* was 1.5%, 1.0%, and 11.3% in healthy replicates A, B, and C, respectively, compared with 36.5%, 46.1%, and 1.2% in diseased replicates A, B, and C. Thus, *Enterobacterales* became one of the dominant orders in diseased rhizoplane samples A and B. In diseased replicate C, by contrast, both *Rhizobiales* and *Enterobacterales* were relatively low, whereas *Burkholderiales*, *Sphingomonadales*, and *Chitinophagales* were more abundant.

**Fig. 1.**
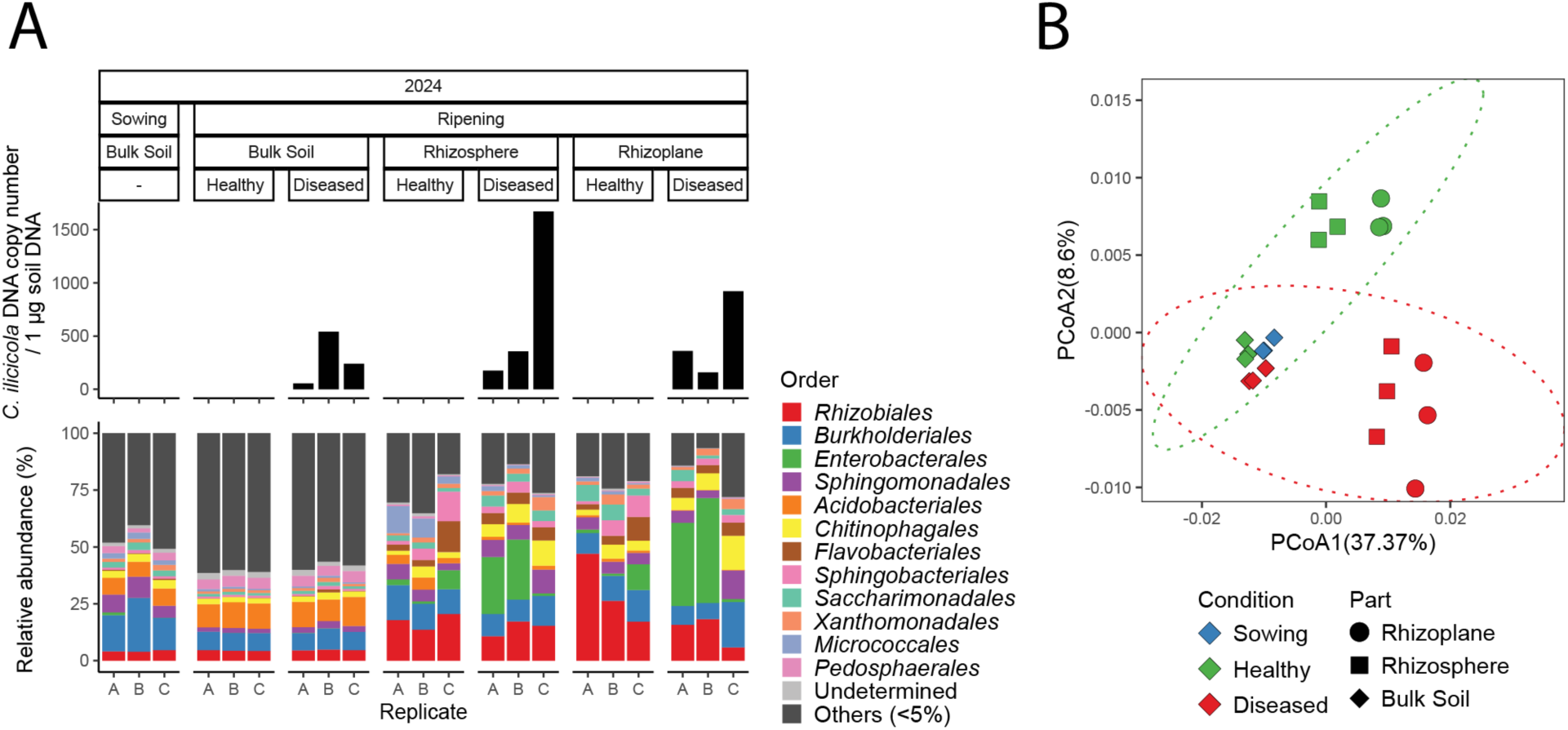
Quantification of *C. ilicicola* and bacterial community structure in soybean soil fractions. (A) The upper annotation boxes indicate, from top to bottom, the sampling year, plant growth stage, soil fraction, and the status of soybean plants. Top: Quantification of *C. ilicicola* DNA copy numbers. The copy number is normalized per 1 µg of total soil DNA. Bottom: Bacterial community composition at the order level based on 16S rRNA gene sequencing. (B) Principal coordinate analysis (PCoA) of weighted UniFrac distances of the bacterial community among all soil fractions collected from the NS4 field in 2024. The dashed line indicates the 90% confidence ellipse.

Principal-coordinate analysis (PCoA) of weighted UniFrac distances further showed that bulk soil communities were similar between the sowing and ripening stages, whereas the rhizosphere and especially the rhizoplane communities of healthy and diseased plants were clearly separated (Fig. 1B).

A similar pattern was observed in the 2022 samples. *C. ilicicola* DNA was detected in all soil fractions from diseased soybean plants except one bulk soil sample. The bulk soils collected at the sowing stage were dominated by *Acidobacteriales*, and *Enterobacterales* increased in the rhizosphere and rhizoplane soils of all three diseased soybean plants, consistent with the 2024 observations (Fig. S2A and B). Together, these results show that soybean RCR is associated with altered bacterial community composition in rhizosphere and rhizoplane soils, particularly in the rhizoplane. Most, but not all, diseased plants showed increased *Enterobacterales* abundance in the rhizosphere and rhizoplane soils, indicating biological heterogeneity among diseased replicates. However, correlation analyses between the quantitative level of *C. ilicicola* DNA and the relative abundance of *Enterobacterales* across soil fractions showed no significant relationship in diseased samples from either 2024 (Pearson’s r = 0.021, P = 0.927) or 2022 (Pearson’s r = 0.093, P = 0.690). These data therefore support an association between disease status and microbiome restructuring, but not a simple linear relationship between fungal load and *Enterobacterales* abundance.

### 3.2. Functional genes related to antibiotic resistance are enriched in the rhizoplane soils with increased *Enterobacterales* abundance

To examine functional shifts in the soil microbiome, we constructed a nonredundant gene catalog comprising 3,537,664 genes from assemblies of 17 shotgun metagenomes, including seven bulk soil samples collected at the sowing stage and 10 samples collected at the ripening stage (two bulk, two rhizosphere, and six rhizoplane samples). Metagenomic reads were mapped to this catalog to compare gene content between healthy and diseased plants. A total of 3,494,533, 3,516,388, and 3,384,625 genes were detected in bulk, rhizosphere, and rhizoplane soils, respectively. Within each fraction, 94.0% (3,285,788/3,494,533) of genes in bulk soils, 95.2% (3,346,316/3,516,388) in rhizosphere soils, and 83.8% (2,837,827/3,384,625) in rhizoplane soils were shared between healthy and diseased conditions (Fig. S3A to C). Bray-Curtis-based comparisons of gene abundance profiles showed that bulk soil functional compositions changed little from sowing to ripening, whereas rhizosphere samples differed between healthy and diseased plants and the separation was most pronounced in rhizoplane samples (Fig. 2A). These results indicate that disease status is associated not only with taxonomic shifts but also with restructuring of microbiome functional profiles.

**Fig. 2.**
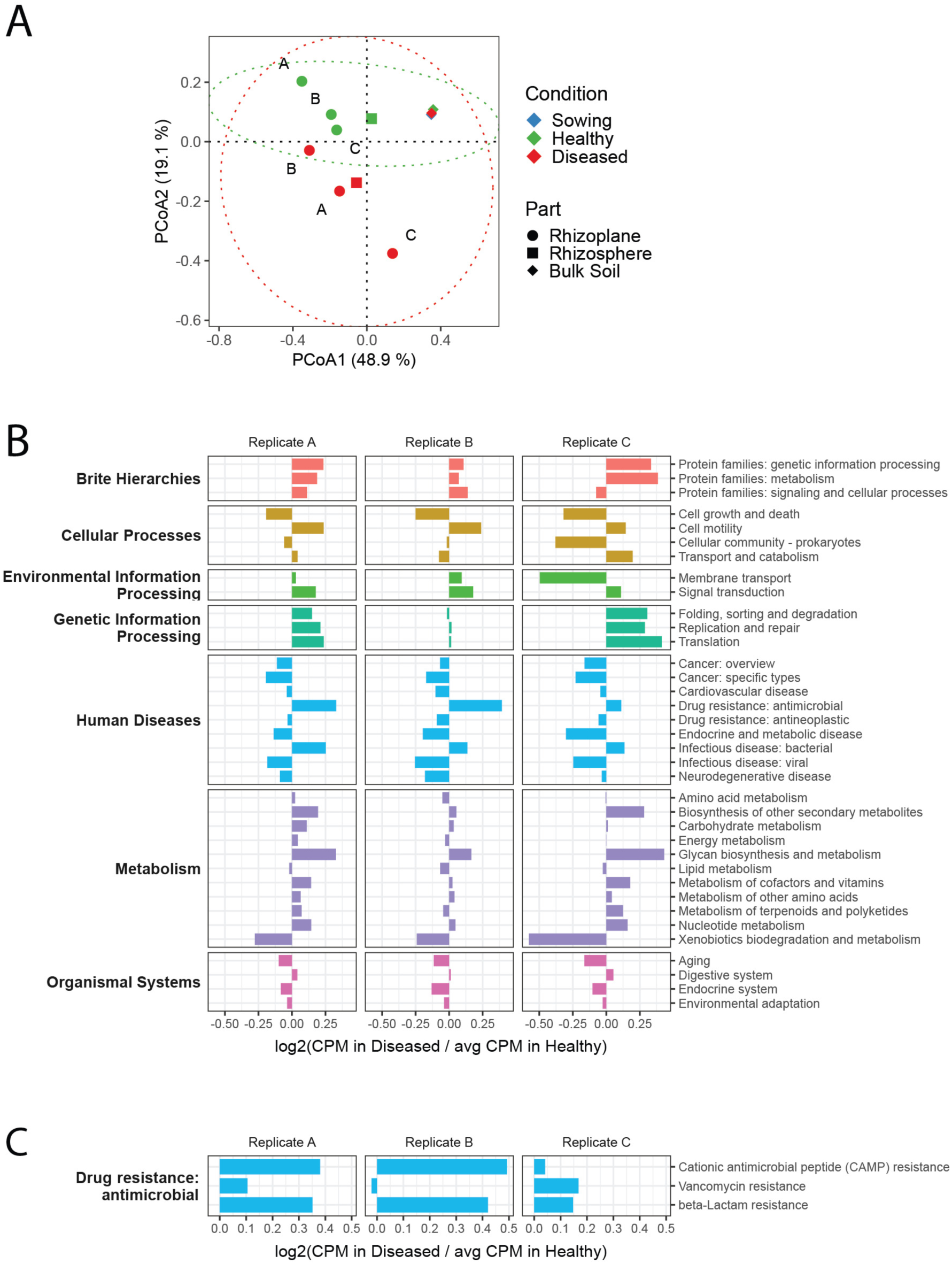
Functional gene composition of soybean soils based on shotgun metagenomic sequencing. (A) PCoA based on Bray-Curtis distances among 11 shotgun metagenome data collected from the NS4 field in 2024. The dashed line indicates the 95% confidence ellipse. (B) Ratios of gene category abundances in diseased rhizoplane soils relative to the mean abundance in healthy rhizoplane soils. Major functional groups are indicated on the left, and detailed gene categories on the right. (C) Ratios of subcategories within “Drug resistance: antimicrobial”, with subcategory names indicated on the right.

We next summarized gene abundances according to KEGG Orthology (KO) categories based on BRITE hierarchies (23) to evaluate functional alterations. For each diseased rhizoplane sample, the abundance of each category was normalized to the mean abundance across the three healthy rhizoplane samples (Fig. 2B).

Among 36 KO categories passing the abundance threshold used for visualization, 15 were consistently increased, and 14 were consistently decreased across diseased rhizoplane samples, indicating broad functional reorganization of the diseased rhizoplane microbiome. The category showing the strongest increase in diseased replicates A and B was “Drug resistance: antimicrobial” which coincided with marked enrichment of *Enterobacterales* in those samples. Within this category, the subcategories “CAMP resistance” and “beta-lactam resistance” were elevated specifically in diseased replicates A and B (Fig. 2C). Relative abundance of the CAMP resistance category increased 1.30-fold and 1.41-fold in diseased replicates A and B, respectively, compared with 1.03-fold in replicate C. Similarly, beta-lactam resistance increased 1.28-fold and 1.34-fold in replicates A and B, compared with 1.11-fold in replicate C. Previous studies have reported that environmental *Enterobacterales* frequently carry beta-lactam resistance genes (24), consistent with a potential contribution of *Enterobacterales* enrichment to this functional shift. The 2022 dataset showed the same overall pattern: samples with higher *Enterobacterales* abundance also showed higher relative abundance of the “Drug resistance: antimicrobial” category, particularly due to increases in the "CAMP resistance” and “beta-lactam resistance” subcategories (Fig. S4A to C).

### 3.3. Single-cell genomics reveals the increase of specific *Enterobacterales* strains in the rhizoplane of RCR

The 16S rRNA gene sequencing and shotgun metagenomics suggested that *Enterobacterales* lineages carrying antimicrobial resistance-related functions were enriched in diseased rhizoplane soils. Because resistance traits can vary substantially among closely related environmental strains (24, 25), we next applied droplet-based single-cell genomics (SAG-gel) to examine *Enterobacterales* populations at higher taxonomic resolution. From pooled rhizoplane soils of healthy and diseased soybean plants, 384 single cells were processed per condition, and the resulting SAG assemblies were evaluated using CheckM (26). Among the recovered SAG assemblies, 18 were classified as high quality, 185 as medium quality, and 504 as low quality. We focused subsequent analyses on the 203 medium- and high-quality SAGs (Table S2).

Among the 203 medium- and high-quality SAGs recovered from rhizoplane soils, *Rhizobiales* was the most abundant order, with 71 SAGs from healthy plants and 49 from diseased plants. Of these 120 *Rhizobiales* SAGs, 106 (88%) were classified as *Bradyrhizobium diazoefficiens*, consistent with the dominance of this symbiotic lineage in soybean root-associated habitats. The second most abundant order was *Enterobacterales*, represented by 51 SAGs from diseased plants and only one SAG from healthy plants. Among these 52 *Enterobacterales* SAGs, 41 (79%) were classified as *Enterobacter asburiae_B* (Table S2). These taxonomic distributions were consistent with the amplicon data and indicate that the SAG-gel workflow successfully recovered genomes from the dominant bacterial lineages present in soybean rhizoplane soils.

To generate a nonredundant genome set, SAGs were clustered with ccSAG to identify redundant assemblies derived from the same strain. This step grouped 86 SAGs into 10 composite genomes, yielding a final nonredundant set of 127 genomes (Table S3). These genomes spanned 17 bacterial orders, with *Rhizobiales* (99 genomes) being most abundant, followed by *Enterobacterales* (7 genomes) (Fig. 3A). Six of the seven nonredundant *Enterobacterales* genomes originated from diseased rhizoplane soils. The *Enterobacterales* genomes had an average completeness of 84.77% (minimum, 63.66%) and an average contamination of 1.39% (maximum, 2.20%) (Table S3). According to GTDB-based classification and comparison with reference genomes, these SAGs corresponded to six *Enterobacterales* species-level lineages (Table S3).

**Fig. 3.**
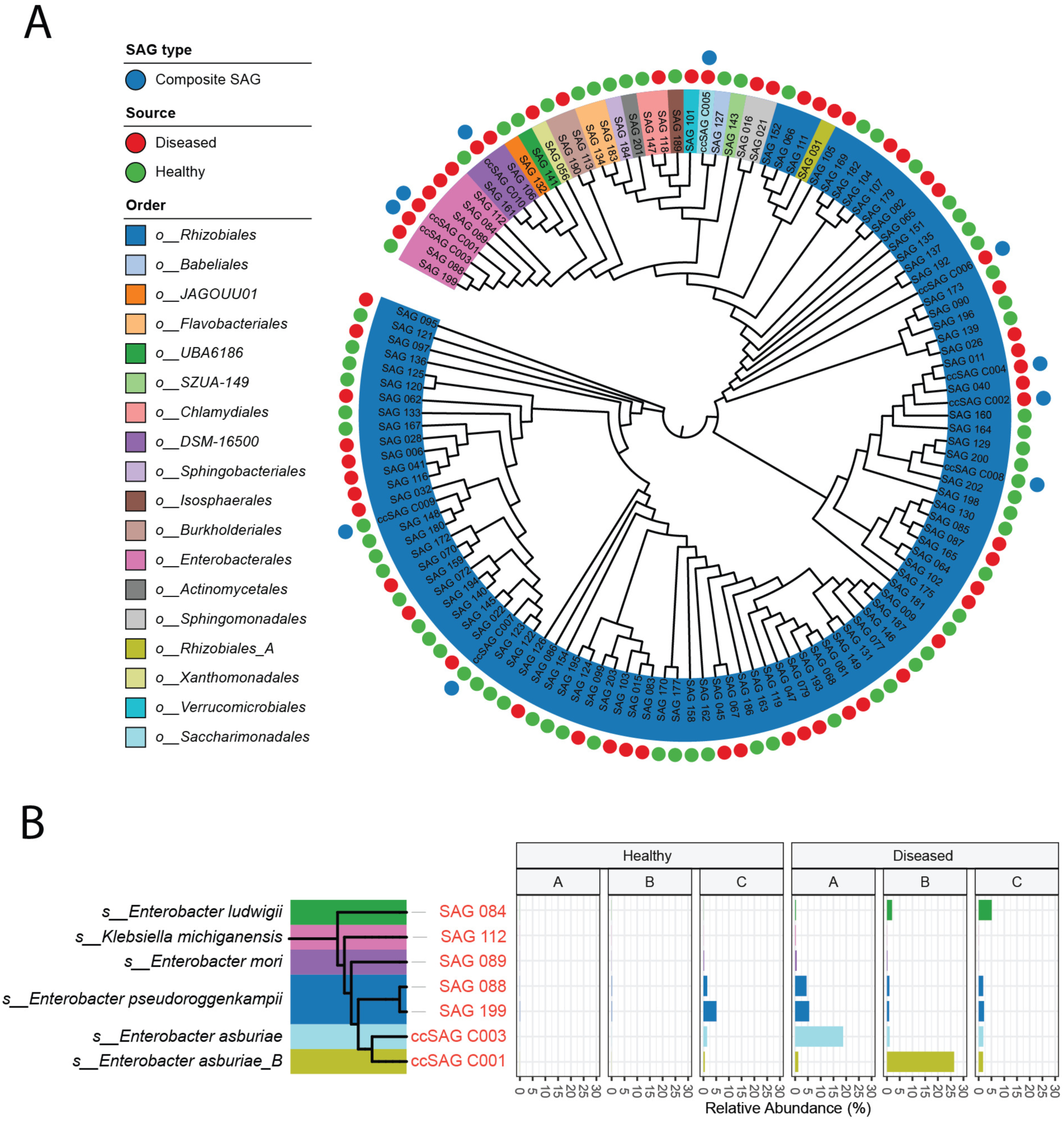
SAGs obtained from rhizoplane soils and their relative abundances in shotgun metagenomes. A) The GTDB-Tk phylogenetic tree of medium or high-quality SAGs obtained from rhizoplane soils. Composite genomes constructed from the same strain SAGs by ccSAG are shown as blue circles. Genomes obtained from rhizoplane soils of diseased soybeans are shown in red circles, and those from healthy soybeans in green circles. The background color of each genome ID indicates its taxonomic annotation at the order level. (B) Relative abundances of the seven *Enterobacterales* SAGs in each rhizoplane soil. The background colors of the GTDB-Tk phylogenetic tree on the left indicate the species of each SAG.

To estimate strain-level abundance in each rhizoplane sample, shotgun metagenomic reads were mapped to the 127 nonredundant genomes. In diseased rhizoplane replicate A, *E. asburiae* and *E. pseudoroggenkampii* together accounted for 92.78% of the total *Enterobacterales* abundance, whereas in diseased replicate B, *E. asburiae_B* alone accounted for 83.86% (Fig. 3B). By contrast, the remaining lineages, including *E. ludwigii*, *Klebsiella michiganensis*, and *E. mori*, each represented less than 6.35% in both diseased replicates A and B. These results indicate that *Enterobacterales* enrichment in diseased rhizoplane soils was driven by a limited number of specific lineages, and that the dominant lineage differed between biological replicates.

### 3.4. *Enterobacterales* strains enriched in the rhizoplane soils of diseased soybean possess antibiotic resistance genes

We next compared the gene repertoires of *Enterobacterales* lineages enriched in diseased rhizoplane soils with those of related reference genomes. For this comparison, we collected 51 high-quality *Enterobacterales* reference genomes from GTDB (27) representing the six species-level lineages detected in our SAG dataset and originating from soil or plant-associated environments. These reference genomes had an average completeness of 99.80% (minimum, 96.98%) and an average contamination of 0.42% (maximum, 1.66%) (Table S4). We then screened each genome for 4,682 plant pathogenicity-related genes associated with plant hosts (*Tracheophyta*) in PHI-base v4 (28), together with four additional genes previously implicated in plant pathogenicity (29, 30). Among the 38 plant pathogenicity-related genes detected in at least one *Enterobacterales* lineage, 25 were shared across all six species-level lineages, indicating that these genes were broadly distributed rather than restricted to lineages enriched in diseased rhizoplane soils (Fig. 4A). These shared genes included genes associated with soft rot-like disease processes and pectin esterase genes implicated in root tissue maceration or root rot pathogenicity (29, 30). Although five genes were lineage specific, none was unique to the lineages enriched in diseased rhizoplane soils.

**Fig. 4.**
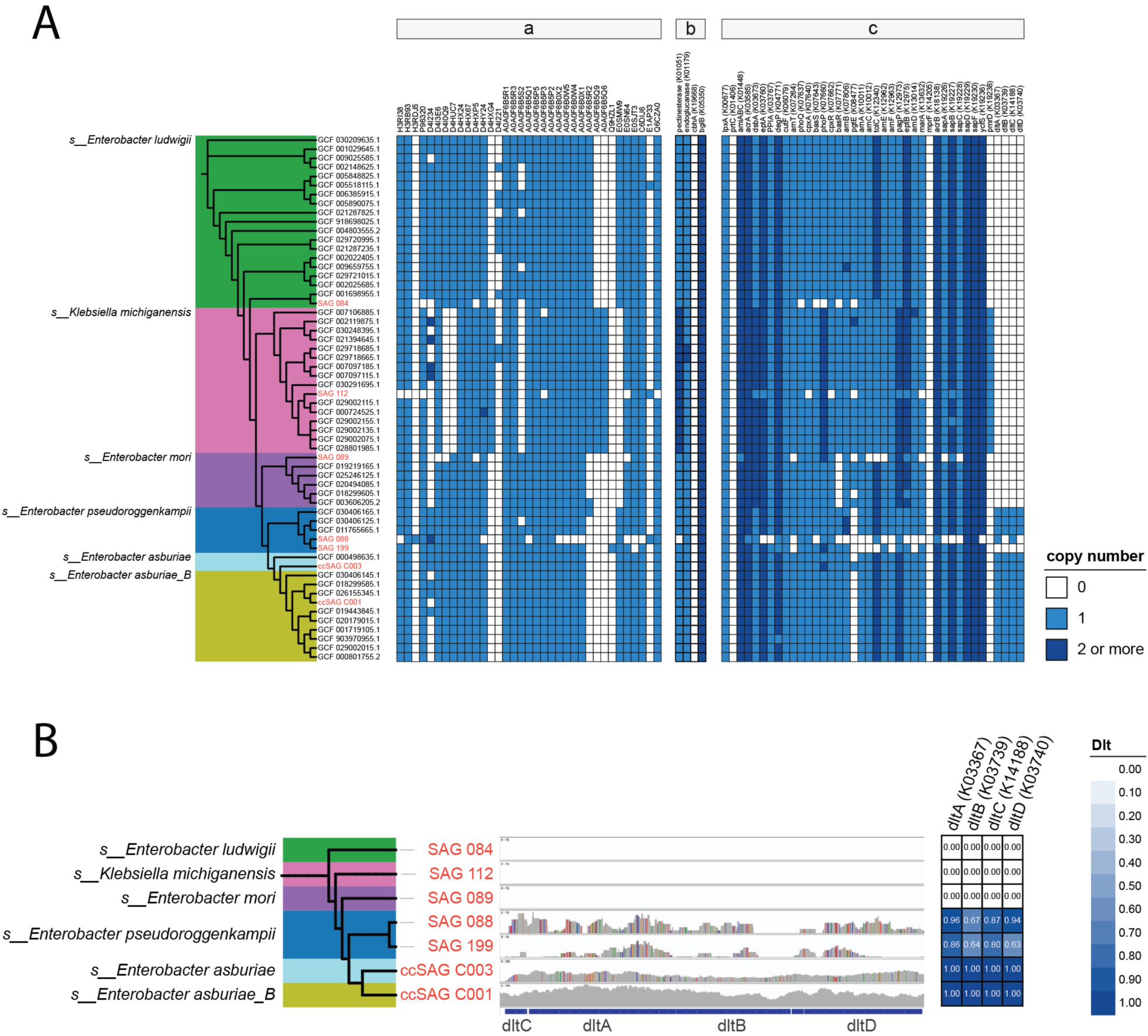
Profiling of pathogenic and CAMP resistance genes in *Enterobacterales* genomes. (A) Possession of plant pathogenic and putative CAMP resistance genes in SAGs and reference genomes of *Enterobacterales*. In the heatmap, white indicates the absence of the gene, light blue indicates a single copy, and dark blue indicates the presence of two or more copies. In the phylogenetic tree, genome IDs in red indicate SAGs obtained from rhizoplane soils in this study, whereas those in black indicate reference genomes.
a: Plant pathogenic genes registered in PHI-base.
b: Additional plant pathogenicity genes reported in the literature.
c: CAMP resistance genes annotated in the KEGG BRITE database. (B) Mapping of sequencing reads from each SAG to the *dlt* operon. The central histogram represents read coverage, and the heatmap on the right shows the proportion of each *dlt* gene region covered by sequencing reads.

We then focused on the “CAMP resistance” functional category, which was enriched in diseased rhizoplane samples A and B. Among the 54 genes assigned to this category, *dlt* genes (*dltA* to *dl*tD) were detected only in the *Enterobacterales* lineages enriched in diseased rhizoplane soils, including *E. asburiae*, *E. pseudoroggenkampii*, and *E. asburiae_B* (Fig. 4A). Because SAG assemblies can be incomplete, we further assessed support at the raw read level. ccSAG_C004 (*E. asburiae*) and ccSAG_C002 (*E. asburiae_B*) contained reads covering 100% of the *dltA*-to-*dltD* reference sequences, and SAG_088 and SAG_199, which belonged to *E. pseudoroggenkampii*, covered 63% to 96% of each *dlt* gene region (Fig. 4B). In contrast, *Enterobacterales* SAGs not enriched in diseased rhizoplane soils showed no detectable read coverage for *dlt* genes. These results suggest that plant pathogenicity-related genes were broadly shared across *Enterobacterales* lineages, whereas *dlt* genes were restricted to the lineages enriched in diseased rhizoplane soils. The *dlt* operon is well known to contribute to CAMP resistance in Gram-positive bacteria and has also been implicated in host-associated survival in some plant pathogenic bacteria (31–33). Although the functional role of *dlt* genes was not tested experimentally in these *Enterobacterales* lineages, its distribution pattern is consistent with a possible role in adaptation to the diseased soybean rhizoplane.

## 4. Discussion

Using 16S rRNA gene sequencing, shotgun metagenomics, and single-cell genomics, we characterized bacterial communities associated with soybean fields affected by red crown rot. 16S rRNA gene sequencing showed that diseased plants had rhizosphere and rhizoplane microbiomes distinct from those of healthy plants, with the clearest separation observed in the rhizoplane (Fig. 1A and B). Across both sampling years (2022 and 2024), most diseased plants showed increased relative abundance of *Enterobacterales* in the rhizoplane. Shotgun metagenomics further indicated that disease-associated rhizoplane communities were functionally shifted toward antimicrobial resistance-related categories, especially CAMP resistance and beta-lactam resistance (Fig. 2 and Fig. S4). Together, these findings support the view that soybean RCR is accompanied by reproducible restructuring of root-associated bacterial communities, particularly at the root surface.

Single-cell genomics provided a strain-resolved context for these community-level patterns. Because soil bacteria are known to exhibit considerable functional diversity even within the same lineage (34–36), single-cell genome sequencing enables direct comparisons of genomic functions among closely related lineages.

Among 203 medium- and high-quality SAGs, *Enterobacterales* were strongly skewed toward diseased rhizoplane soils, and read recruitment to the nonredundant genome set showed that only a small subset of lineages accounted for most of the *Enterobacterales* abundance in diseased samples. Comparison with related reference genomes showed that plant pathogenicity-related genes were widespread across *Enterobacterales* lineages, whereas *dlt* genes, putative CAMP resistance genes, were restricted to the lineages enriched in diseased rhizoplane soils.

Although the shared pathogenicity-related genes may still contribute to colonization or tissue deterioration after establishment, they did not distinguish the disease-associated lineages in our dataset. This contrast suggests that, within this dataset, putative tolerance to host-derived antimicrobial peptides distinguishes disease-associated *Enterobacterales* lineages more clearly than the mere presence of generic pathogenicity-related genes. To our knowledge, this is the first genome-resolved study to examine bacterial community structure and strain-level functional variation in the rhizosphere and rhizoplane of soybean plants affected by RCR.

Plants secrete AMPs and other defense-related compounds from their roots in response to microbial challenge (15). In soybean, CAMPs such as defensins and glycinin-derived peptides have been reported to inhibit the growth or infection ability of certain pathogens (37, 38). Our data are therefore consistent with a tripartite model in which fungal infection alters the soybean root environment, host defense responses impose selective pressure, and only a subset of bacterial lineages with suitable gene repertoires can proliferate. In this framework, *Enterobacterales* lineages carrying *dlt* genes may be better able to persist, or even expand opportunistically, in the chemically altered rhizoplane of diseased plants. At the same time, the present study does not demonstrate that these bacteria initiate disease or directly exacerbate symptoms. Rather, the data support an association between disease status and enrichment of specific lineages with gene repertoires potentially suited to survival in the diseased root environment. This interpretation is also compatible with previous reports showing that multiple microbial pathogens or opportunists often coexist in soybean root diseases (39).

Several limitations should be acknowledged. First, *Enterobacterales* enrichment was not observed in every diseased plant; for example, diseased replicate C in 2024 did not show an increase, whereas healthy replicate C still contained 11.3% *Enterobacterales*. This heterogeneity suggests that *Enterobacterales* expansion is not a defining feature of all RCR-affected plants and may depend on disease stage, local microenvironmental conditions, or both. Second, no clear correlation was observed between *Enterobacterales* abundance and *C. ilicicola* DNA level across soil fractions, indicating that *Enterobacterales* enrichment is neither necessary nor sufficient for RCR symptoms. Third, some shotgun libraries and all SAGs were based on pooled material, which limits replicate-level inference for part of the functional and strain-resolved analyses. Finally, this study captured a single time point at the ripening stage and therefore cannot resolve temporal order or causality. It remains possible that *Enterobacterales* become prominent only in a subset of diseased roots or at particular stages after host defense responses intensify. Even with these limitations, the present dataset provides a useful framework for identifying bacterial lineages and gene functions consistently associated with diseased soybean rhizoplanes. Because no additional experiments are included here, we intentionally interpret these bacterial features as disease-associated rather than disease-causing. In this form, the work provides a genome-resolved foundation for future studies on diagnostic markers, microbial interactions, and disease-suppressive strategies in soybean production systems. Future integration of host-centered analyses, including soybean transcriptomics, root exudate profiling, and root histology, with genome-resolved microbiome data may further clarify the progression of RCR and the context in which *Enterobacterales* become enriched (40, 41).

## 5. Method

### 5.1. Soybean Sampling and Soil DNA Extraction

In June, soybean (cultivar: Ryuhou) was sown in experimental fields located in Daisen City, Akita Prefecture, and bulk soils were collected in triplicate at the sowing stage. Sampling was conducted in 2022 and 2024: in 2022, soils were obtained from field NS1, while in 2024, bulk soils were collected from seven fields (NS1-NS7) (Table S5). All fields were converted paddy fields with soils classified as Fluvisols according to the World Reference Base for Soil Resources (42). The maximum distance between fields was 838 m, observed between NS1 and NS5.

Fifteen weeks after sowing, poorly growing soybeans exhibiting root rot symptoms were identified at NS1 in 2022 and at NS4 in 2024. From each of these fields, three poorly growing and three healthy soybeans were uprooted, and roots with adhering soil were collected. In addition, bulk soils were sampled at a distance of more than 15 cm from the plant base and at a depth of approximately 10 cm. This sampling time corresponded to the ripening stage. Rhizosphere and rhizoplane soils were fractionated from the soybean samples following a previously described protocol (43). As a result, a total of 60 soil fractions were obtained at the sowing and ripening stages. For soybean samples collected in 2024, a portion of the rhizoplane soil was stored at 4℃ for subsequent single-cell genome amplification. Microbial DNA was extracted from each soil using the Extrap Soil DNA Kit Plus ver. 2 (Biodynamics Laboratory, Tokyo, Japan). DNA was eluted in 100 µL of buffer for the bulk soils, and in 50 µL for the rhizosphere and rhizoplane soils. After elution, an equal volume of 10% PVPP solution was added and mixed. The mixture was then centrifuged at 14,000 × g for 3 min at 20℃, and 30 µL of the supernatant was recovered. DNA concentrations were measured using a Qubit Fluorometer (Thermo Fisher Scientific, Waltham, MA, USA).

### 5.2. Quantification of *C. ilicicola* DNA by qPCR

Quantitative PCR was performed to determine the amount of *C. ilicicola* DNA in all 60 soil DNA samples. Previously reported primers targeting the IGS region of *C. ilicicola* were used: CiIGSF (forward, 5’-TCCATTGCCTCTATTTATCCTGC-3’), CiIGSR (reverse, 5’-GCGTAAAGATTTTCCAACCCG-3’), and probe CiPro (5’-ACCACAGCACAACACGCACAAC-3’) (10). The probe was labeled with fluorescein amidite (FAM) at the 5’ end, an Iowa Black fluorescent quencher (IBFQ) at the 3’ end, and a ZEN quencher internally. *C. ilicicola* DNA was extracted from *C. ilicicola* strain AP-12 (44) cultured on PDA medium at 25℃ using the Wizard® Genomic DNA Purification Kit (Promega, Madison, WI, USA) according to the manufacturer’s instructions. The concentration of extracted DNA was evaluated with a Qubit Fluorometer (Thermo Fisher Scientific) and used as a standard for qPCR. A standard curve was generated using a 10-fold serial dilution of *C. ilicicola* DNA (5×10^−2^, 5×10^−3^, 5×10^−4^, 5×10^−5^ ng/µL). Quantitative values that did not meet the threshold of 6.99 × 10⁻¹ copies per µL of input DNA were regarded as below the detection limit. qPCR was conducted using PrimeTime® Gene Expression Master Mix (Integrated DNA Technologies, Coralville, IA, USA) according to the manufacturer’s instructions. For each soil DNA sample, 1 µL was analyzed in triplicate, and the amount of *C. ilicicola* DNA was calculated. Mean values from triplicate measurements were normalized to the input DNA concentration, yielding the quantity of *C. ilicicola* DNA per 1 µg of extracted DNA. The Copy number was estimated based on the molecular weight of the *C. ilicicola* reference genome (NCBI Assembly accession: GCA_020809705.1) (4.31×10^10^ g/mol). Data visualization was performed using the R package ggplot2 (45, 46).

### 5.3. 16S rRNA Gene Sequencing and Analysis

Bacterial community analysis was performed on all 60 soil DNA samples. Library preparation followed a previously described protocol (43) targeting the V3-V4 hypervariable region of the 16S rRNA gene. Amplicons obtained in 2022 were sequenced using the MiSeq Reagent Kit v3 on the Illumina MiSeq platform (2 × 300 bp; Illumina, San Diego, CA, USA) according to the manufacturer’s instructions.

Amplicons from 2024 were sequenced using the NextSeq P1 600 Cycle Reagent Kit on the Illumina NextSeq 2000 platform (Illumina). Following the same protocol (43), primer sequences were removed, an ASV count table was generated, taxonomy was assigned to ASVs, and eukaryotic ASVs were removed. Community composition was analyzed using nyankomicro (47). PCoA based on weighted UniFrac distance was conducted in R (version 4.1.2) using the Bioconductor package phyloseq (48) and MicrobiotaProcess (49). Correlation analyses between the quantitative levels of *C. ilicicola* DNA and the relative abundance of *Enterobacterales* were performed using the cor.test function in R (46).

### 5.4. Shotgun Metagenome Sequencing and Analysis

A total of rhizoplane, rhizosphere, and bulk soil samples collected in 2022 and 2024 were used for library preparation. For each rhizoplane soil, 7 µL of extracted DNA was independently subjected to library preparation. In contrast, DNA from triplicate rhizosphere and bulk soils was pooled at equal DNA concentrations, and 7 µL of the pooled DNA was used for subsequent analysis. For shotgun metagenome sequencing, DNA from bulk soil at the sowing stage and from bulk and rhizosphere soils at the ripening stage in 2022, as well as from bulk and rhizosphere soils at the ripening stage in 2024, was sequenced by bitBiome, Inc. (Tokyo, Japan) using the DNBSEQ platform (MGI Tech Co., Shenzhen, China) to generate 150 bp paired-end reads. DNA from rhizoplane soil at the ripening stage in 2022 and from bulk soil at the sowing stage and rhizoplane soil at the ripening stage in 2024 was sequenced in our laboratory. DNA libraries were prepared using the QIAseq FX DNA Library Kit (QIAGEN, Hilden, Germany) and sequenced on the Illumina NextSeq 2000 platform (Illumina) with the NextSeq 2000 P3 XLEAP-SBS Reagent Kit (300 cycles). A total of 17 shotgun metagenomic datasets were obtained. Reads were quality-controlled using bbduk.sh and bbmap.sh (50), followed by assembly with SPAdes v3.15.2 (51).

Gene prediction was conducted with Prokka v1.14.6 (52), and functional annotation was assigned using EggNOG-mapper v2.1.6 (53). MMseqs2 v17.b804f (54) was used to construct a nonredundant gene catalog. Reads were mapped to the catalog with CoverM v0.7.0 (55) to generate counts per million (CPM) tables. PCoA based on Bray-Curtis distance was performed using the pcoa function in the R package ape (56). KEGG BRITE database entries (as of March 4, 2024) were used to aggregate CPM values by BRITE hierarchies (23). For each category and each subcategory within "Drug resistance: antimicrobial", the ratio of CPM values in each rhizoplane soil sample of diseased soybeans against the average CPM of the three healthy soybean rhizoplane soils was calculated. Only categories with a total CPM of at least 15,000 across the six rhizoplane soils were visualized using ggplot2. (45, 46).

### 5.5. Single-Cell Genome Sequencing Using the SAG-gel Method

Rhizoplane soils collected in 2024 from three healthy and three diseased soybean plants were separately pooled into a healthy sample set and a diseased sample set. Following a previously described protocol (43), pooled soils were subjected to single-cell genome sequencing using the SAG-gel method (17, 18). Following the same protocol (43), low-quality reads were removed, assemblies were generated, and genome completeness and contamination were evaluated. Genome quality categories were assigned as described previously (57). Taxonomic classification was conducted using GTDB-Tk v2.4.1 (58) with default options against the Release 226 database. To identify strains, SAGs were clustered using ccSAG v2.1.1 (59) (options: -criteria-ani 99 -criteria-marker 0.99). Redundant SAGs were merged using ccSAG clean (options: -total-completeness 200 -total-contamination 10), yielding nonredundant SAGs. A phylogenetic tree of the nonredundant SAGs was constructed using GTDB-Tk infer (58) and visualized with iTOL (60). Shotgun metagenome reads from rhizoplane soil of healthy and diseased soybean in 2024 were mapped to the nonredundant SAGs using CoverM v0.7.0 genome mode (55) (options: -min-read-percent-identity 95 -min-read-aligned-percent 75 -methods tpm -mapper bwa-mem). The relative abundance of each SAG was calculated as its proportion of the total transcripts per million (TPM) count within each sample and visualized using ggplot2 (45, 46).

### 5.6. Gene Content Analysis of *Enterobacterales* Genomes

To investigate general gene possession patterns among *Enterobacterales*, high-quality genomes annotated as isolated from soil or plants in GTDB (27) were collected. Gene prediction was performed with Prokka v1.14.6 (52), and KEGG Orthology annotation was assigned using EggNOG v2.1.6 (53). Plant pathogenicity genes were identified using PHI-base v4 (28), and Additional plant pathogenicity genes were obtained from the literature (29, 30). The collected pathogenic genes were searched against both reference genomes and SAGs of *Enterobacterales* using BLAST+ (61), with thresholds of ≥ 90% sequence identity and ≥ 90% alignment coverage. KEGG Orthologies corresponding to CAMP resistance genes were identified based on KEGG BRITE database (23), and their presence in each genome was determined according to the KEGG Orthology annotations assigned to the genomes (23). Gene presence was visualized as heatmaps in iTOL (60). To assess read-level coverage, a contig containing the *dlt* gene from ccSAG C001 (*E. asburiae_B*) was used as a reference. Reads from each SAG were mapped to this contig using CoverM v0.7.0 (55) contig mode, and BAM files were generated, sorted, and indexed with Samtools v1.21 (62). Visualization was performed using IGV (63).

## 7. Acknowledgments

This work was supported by Cabinet Office, Government of Japan, Moonshot R&D Program for Agriculture, Forestry and Fisheries (JPJ009237, funding agency: Bio-oriented Technology Research Advancement Institution). We appreciate the helpful discussions and insightful comments provided by Ryota Wagatsuma. We also thank Chikako Sakanashi, Kaori Aikawa, Yuki Ohnishi, and Koki Kashiwagi for their assistance with sampling and experimental work.

## 8. Supplementary Materials

Supplementary Figures – Supplementary_Figures.pdf

Figures S1 to S4

Supplementary Tables – Supplementary_Tables.xlsx

Table S1 – Quantitative results of qPCR targeting *C. ilicicola* DNA (in triplicate).

Table S2 – Metadata of SAGs with medium or higher quality.

Table S3 – Metadata of non-redundant SAGs with medium or higher quality.

Table S4 – Metadata of reference genomes obtained from the GTDB.

Table S5 – Coordinates of the field where sampling was conducted.

## 9. Data Availability

All raw sequencing data generated in this study, including 16S rRNA amplicon, shotgun metagenomic, and single-cell genomic datasets, have been deposited in the NCBI database under BioProject accession number PRJNA1338991.

